# Oral Streptococci *S. anginosus* and *S. mitis* induce distinct morphological, inflammatory, and metabolic signatures in macrophages

**DOI:** 10.1101/2023.08.28.555100

**Authors:** Sangeetha Senthil Kumar, Venugopal Gunda, Dakota M. Reinartz, Kelvin Pond, Paul Victor Santiago Raj, Michael D. L. Johnson, Justin E. Wilson

**Author notes:** **Corresponding Authors** Justin E Wilson, Ph.D. Michael D. L Johnson, Ph.D. Department of Immunobiology 1656 E. Mabel Street, PO Box 245221, Tucson, AZ 85724-5221.

## Abstract

Oral streptococci are the pioneer colonizer and structural architect of the complex oral biofilm. Disruption of this architectural framework causes oral dysbiosis associated with various clinical conditions, including dental caries, gingivitis, and oral cancer. Among the genus *Streptococcus*, *S. anginosus* is associated with esophageal, gastric, and pharyngeal cancer tissues, while *S. mitis* is correlated with oral cancer. However, no study has investigated mechanistic links between these *Streptococcus species* and cancer-related inflammatory responses. To explore the underlying involvement of *S. anginosus and S. mitis* in inflammation-associated cancer development, we investigated the innate immune response elicited by *S. anginosus* versus *S. mitis* using the RAW264.7 macrophage cell line. Compared to untreated or *S. mitis* infected macrophages, *S. anginosus* infected macrophages exhibited a robust proinflammatory response characterized by significantly increased levels of inflammatory cytokines and mediators, including TNF, IL-6, IL-1β, NOS2, and COX2, accompanied by enhanced NF-κB activation. Mitostress analysis revealed an increased extracellular acidification rate in macrophages infected with *S. anginosus* compared to *S. mitis.* Further, macrophages infected with *S. anginosus* for 6h displayed upregulated aconitate decarboxylase, which catalyzes itaconate production. In contrast, no significant alterations were observed in succinate dehydrogenase that converts succinate to fumarate. At 24h, *S. anginosus* induced significant shifts in succinate and itaconate, emphasizing a unique macrophage metabolic profile, and an augmented inflammatory response in response to *S. anginosus.* This study underscores the capacity of *S. anginosus* to elicit a robust proinflammatory response in macrophages and opens new avenues of secretory immune metabolites in response to oral streptococci.

**Importance:** The surge in head and neck cancer cases among individuals devoid of typical risk factors such as HPV infection, tobacco and alcohol use sparks an argumentative discussion around the emerging role of oral microbiota as a novel risk factor in oral squamous cell carcinoma (OSCC). While substantial research has dissected the gut microbiome’s influence on physiology, the oral microbiome, notably oral streptococci, a gatekeeper of systemic health, has been underappreciated in mucosal immunopathogenesis. *S. anginosus,* a viridans streptococci group, has been linked to abscess formation and an elevated presence in esophageal cancer and OSCC. The current study aims to probe the innate immune response to *S. anginosus* compared to the early colonizer *S. mitis* as an initial ride towards understanding the impact of distinct oral *Streptococcus* species on the host immune response in the progression of OSCC.

## Introduction

Owing to its thermally and humidly conducive environment, coupled with nutrient circulation facilitated by saliva, the oral cavity accommodates a broad spectrum of microbial species. It encompasses approximately 700 distinct taxa, exhibiting varied compositions across discrete anatomical locales, including the tongue, palate, teeth, and cheeks (1). Oral streptococci, also called viridans group streptococci (VGS), are the pioneer colonizers and the dominant colonizers of oral biofilm. These cocci belong to the Gram-positive category and are catalase-negative facultative anaerobes; most are alpha-hemolytic while some exhibit β or no hemolysis on blood agar (2). Being early colonizers of the oral cavity *Streptococcus* species like *S. mitis, S. sanguinus, S. gordonii, S. oralis*, and *S. salivarius* (3) play a significant role in maintaining oral homeostasis by establishing stable multispecies dental biofilm (4). Ecological pressures imposed by inadequate oral hygiene, dietary changes, or antibiotic usage can reduce this dominant population and disturb biofilm homeostasis resulting in oral dysbiosis. Oral dysbiosis culminates in several oral diseases and infections, such as periodontitis, gingivitis, and oral ulcers, that might drive mucosal breach and immunopathogenesis of various chronic diseases, including cancer (5). The *Streptococcus anginosus* group (SAG), formerly known as the *S. milleri* group, refers to a subgroup of the VGS composed of three species, namely *S. anginosus*, *S. constellatus*, and *S. intermedius*. Although SAG bacteria were first detected in dental abscesses, leading to oral infections, their pathogenicity has extended beyond oral cavities, and they are now increasingly identified as causative agents of infections in various anatomical locations, exhibiting species-specific clinical characteristics (6,7). Systemic *S. anginosus* infections have been reported in patients with esophageal, gastric, and oral cancer (8,9). Furthermore, *S. anginosus* is an especially relevant marker of head, neck, and esophageal cancers and is more common in oral squamous cell carcinoma. (10,11). *S. mitis*, another commensal and one of the early colonizers of the oral cavity, is also reported to be significantly elevated in the saliva of patients with OSCC with a diagnostic sensitivity and specificity of 80% and 82%, respectively (12). In addition, *S. mitis* is reportedly involved in bacteremia and endocarditis (13). Studies have shown that the abundance of saccharolytic and aciduric species, including *S. mitis*, is considerably higher on tumor surfaces than on non-tumor tissue. Although, a higher prevalence of *S. mitis* and *S. anginosus* has been observed in cancerous esophageal tissue (14). The potential implications of this microbial enrichment on the carcinogenic process remain unclear and warrant further investigation (15). Understanding the host response to these bacteria may provide insights to identify the role of these streptococci species in carcinogenesis.

Macrophages act as immune system sentinels, detecting microbes via Pattern Recognition Receptors (PRRs) and triggering the production of reactive molecules and cytokines, which are crucial for our immune response (16). Macrophages exhibit varied behavior between M1 and M2 phenotypes. M1 macrophages excel in generating reactive molecules and specific signals, bolstering pathogen defense. On the other hand, M2 macrophages are activated by type 2 cytokines such as IL4 and IL13, and they assist in the reduction of inflammation, as well as wound healing, and response to helminth infections and allergies. M1 macrophages are identified by unique surface markers like CD80, CD86, and CD16/32. The distinctive cytokines produced by M1 include Interleukin-1β (IL-1β), Interleukin-6 (IL-6), Tumor Necrosis Factor (TNF), Interferon-gamma (IFN-γ), Interleukin-12 (IL-12), and Interleukin-23 (IL-23). On the other hand, M2 macrophages express markers such as arginase-1, mannose receptor (CD206), anti-inflammatory cytokine IL-10, and chemokines CCL17 and CCL22 (17). In bacterial infections, macrophages usually increase the expression of genes related to M1 polarization. This pattern signifies a robust activation that acts as a general alarm against bacteria. Interestingly, these genes are often triggered regardless of the specific bacterial species and offer protection during acute infectious diseases. (18).

Despite being recognized for their association with dental infections and subacute bacterial endocarditis, the presence of VGS in various host body fluids and tissues, notably the heightened colonization of *S. anginosus* and *S. mitis* within squamous cell carcinoma, remains insufficiently explored in terms of their complex interplay with the host. In this study, we investigated the morphological, inflammatory, and metabolic attributes of the macrophage cell line RAW264.7 upon exposure to S. *anginosus* and *S. mitis* infections. The key findings indicate that *S. anginosus* evokes a proinflammatory response comparable to lipopolysaccharide (LPS) in 6h post-infection. However, it demonstrates a distinct extracellular metabolite profile from LPS alone treated macrophages in 24h post-infection. On the other hand, *S. mitis* prompts a subdued 6h inflammatory reaction compared to LPS and *S. anginosus* infection while releasing some immune-regulatory metabolites in 24h post-infection like LPS treatment. Our data indicate that these *Streptococcus* species evoke discrete immune and immunometabolism responses, potentially influencing their inclination to function as either commensal microorganisms or virulent pathogens.

## Materials and Methods

### Bacteria, Chemicals, and Reagents

*Streptococcus anginosus* (ATCC 33397) and *Streptococcus mitis* (ATCC 49456) bacterial strains, along with RAW 264.7 macrophages (TIB-71™), were procured from the American Type Culture Collection (Manassas, VA, USA). BD BACTO™ Brain Heart Infusion and tryptic soy agar were acquired from Fischer Scientific, PA, USA. The culture medium used was Dulbecco’s Modified Eagle’s Medium (DMEM) supplemented with glutamine and sodium pyruvate (Corning Life Science, NY, USA). For gene expression analysis of cytokines and inflammatory mediators—namely, TNF (Mm0043258), IL6 (Mm00446190), IL1b (Mm00434228), Cox2/ptgs2 (Mm00478374), and Nos2 (Mm0040502) Taqman probes were sourced from Thermofisher Scientific, Inc. (MA, USA). The Mouse TNFα, IL-6, and IL1β DuoSet ELISA kits were procured from R&D Systems (MN, USA). For protein analysis, anti-rabbit monoclonal antibodies for p65 (#8242S), phospho-p65 (#3033S), total IKBα (#4812S), phospho-IKBα (#2859S), NLRP3 (#15101S), COX2 (#12282), and Irg1/ACOD1 (17805S) were purchased from Cell Signaling Technology (Danvers, MA, USA). The anti-SDHA antibody (ab14715) and anti-iNOS antibody were procured from Abcam (Waltham, MA, USA). All other chemicals were obtained from Sigma Chemicals (St. Louis, MO, USA).

### Colony Formation Unit (CFU) Assay

CFU assay was performed following the protocol of Menghani et al.(19). A loop of *S. anginosus* and *S. mitis* glycerol stocks was inoculated on blood agar plates and incubated at 37°C in a CO2 incubator overnight. Following this incubation, a loop of the resultant culture was removed from the blood agar plate and added to 10 mL of pre-warmed brain heart infusion broth, and the growth was monitored until the OD600 became 0.1. Upon achieving the desired OD600, serial dilutions of the broth culture were prepared from 100 µl of the stock culture in 96 well plates, with 10 µl of each dilution plated on the blood agar (n=6). The plates were then incubated for 24 hours in a CO2 incubator at 37°C. The colony-forming units (CFU) were then enumerated to determine the viable cell count (Supplemental Figure. 1).

### Bacteria and macrophage coculture

On day 1, RAW 264.7 macrophages were grown as an adherent monolayer culture in 1X DMEM supplemented with 10% Fetal Bovine Serum (FBS) and 5% penicillin-streptomycin. Cells were maintained in humidified air containing 5% CO2 at 37°C overnight. On day 2, the media was removed, cells were washed with PBS, and DMEM medium with FBS devoid of antibiotics were added. Meanwhile, On day 1, a loop of glycerol stock obtained from S. anginosus and S. mitis was streaked on a blood agar plate and incubated at 37°C in a CO2 incubator for overnight growth. On day 2, a loop of bacteria grown overnight on the blood agar plate was scrapped and inoculated in brain heart infusion (BHI) broth at 37°C with 5% CO2 and allowed to grow until an OD600 of 0.1 Then the appropriate amount of culture corresponding to the desired multiplicity of infection (MOI) was pipetted and pelleted by centrifuging @ 4000 rpm for 10 min at 4°C. The bacterial pellet was then resuspended in 1ml PBS, and 100ul of the resuspension was used to infect the macrophages with an MOI of 10.

### Cell Fixation in Culture Plates for macrophage growth and morphology

The protocol for fixing adherent cells was adapted by Martis et al. (20) with slight modifications. In brief, RAW264.7 cells were grown in 24 well plates treated with LPS, *S. anginosus*, or *S. mitis* for 6h or 24h with the MOI of 1, 10, or 50. After treatment, the media was removed, and the cells were washed twice with phosphate-buffered saline (PBS). Then, 0.5 mL of 4% paraformaldehyde (PFA) was added to the cells and incubated for 15-20 minutes at room temperature. The PFA was aspirated, and the cells were washed twice with ice-cold PBS with a 5-minute wait time between each wash. Then, 1 mL of PBS was added, and the plate was wrapped and stored at 4°C until the images were captured in an ECHO Revolve D2224.

### Crystal Violet Staining

Crystal violet staining was performed according to the methods of Zinser et al. (21) and Feoktistova et al. (22). The cells were fixed as described above and stored in PBS at 4°C. On the day of staining, the PBS was removed, and 200 μL of 0.25% crystal violet was added for 30 minutes. The stain was removed, and the plate was washed in running tap water, immersing it in an appropriate container until it became clear. The plate was then blotted on a paper towel, air-dried for 15 minutes, and the images were captured at 20x an ECHO Revolve D2224.

### Giemsa Staining

Giemsa staining was carried out by adopting the method used to quantify phagocytosis by Nicola et al. (23). RAW264.7 cell lines were plated in 96 well plates at the seeding density 5×10^4^. The next day, the media was removed, and the cells were infected with *S. anginosus* or *S. mitis* with DMEM media containing no antibiotics at an MOI of 10. The cells were centrifuged at 24°C for 10 min at 500g. The plate was incubated at 37°C in a 5% CO2 incubator for 3h. After the 3-hour incubation, the cells were fixed with 200 ul of ice-cold methanol for 30 min. Methanol was removed, and each well was washed 2X with 200ul of PBS followed by the 100ul of Giemsa working solution, wrapped, and kept at 4°C overnight. After overnight incubation, the staining solution was aspirated, and the wells were washed gently with PBS. Cells were observed, and the images were captured using a Biotek citation at 40X magnification. At least 300 cells per condition were quantified using NIS-Elements image analysis software. The area of both the cytoplasm and the nucleus was quantified manually to obtain the nucleus-to-cytoplasmic ratio.

### RNA isolation and qPCR

Following 6h post-infection, media was removed from the macrophages, which were then washed with PBS, and 750ul of Trizol was added per well of 6 well plate. The plate was sealed and stored at −80°C until the RNA isolation. RNA extraction was carried out by adopting the phenol/chloroform method published in Toni et al (24). In brief, Cells were scrapped in Trizol, mixed, and transferred to Eppendorf tubes, followed by the addition of chloroform, centrifuged at 12000xg for 15 min at 4°C.The upper phase was transferred and precipitated with isopropanol. Pellets were washed in ethanol, dried, resuspended in 20 μl nuclease-free water, and quantified using nanodrop. cDNA synthesis was performed iScript kit (Biorad, Hercules, CA) in a final volume of 20 uL with the following temperature and time points: 25°C for 5 min, 46°C for 20 min, and 95°C for 1 min. qPCR analysis was carried out in 96 well plates using Quantstudio 3 (Eppendorf). Amplification was carried out at 95°C for 15 min and 50 cycles at 95°C for 15s, 55°C for 30s, and 72°C for 60s, according to Applied Biosystems. The threshold cycle (CT), which indicates the relative abundance of the transcript, was used to quantify the gene expression using the delta Ct method normalized to GAPDH.

### Enzyme-Linked Immunosorbent Assay

ELISA for TNFα (DY410-05), IL6(Dy-406-05), and IL1β(DY401-05) were performed using the R&D systems kit per manufacturer’s instructions, and the absorbance was read using an ELISA reader (BIO-RAD) at 450 nm and 570 nm dual filters.

### Western blot

Western blot was done by adapting the protocol published by Wilson et al. (25) with minor modifications. In brief, total protein lysates were generated from the macrophage cultures by washing cells in PBS and then lysing the cells in ice-cold RIPA buffer containing complete protease inhibitor and phosSTOP. The cell lysate was cleared of insoluble material by centrifugation at 14000g for 10 min @ 4°C followed by separation on SDS-polyacrylamide gel (gel % according to the protein size) and transfer to nitrocellulose membranes (Thermoscientific, USA). After blocking with 10% nonfat dried milk in TBST (Tris buffer saline tween-20) for 1 hour, membranes were incubated with primary antibody overnight at 4°C on a gel rocker in 5% nonfat dried milk. Blots were washed with TBST and incubated with a secondary antibody for 1.30h at room temperature. Membranes were rewashed three times with TBST. The membranes were incubated with ECL reagent in the dark. The membrane blots were exposed to X-ray film and processed using an automatic film processor (OPTIMAX). Band intensity was semi-quantitated using densitometric analysis (ImageJ software).

### Metabolic flux asay

Oxygen consumption (OCR) and extracellular acidification (ECAR) rates were measured using SeaHorse Extracellular Flux (XFe96) analyzer (Agilent Bioscience, US). After optimizing the seeding density through a series of experiments, RAW 264.7 cells were plated in a seahorse microculture plate (1.4× 104 cells/well) in DMEM with FBS and antibiotics and incubated overnight for the cells to adhere. The next day, the sensor and cartridge were hydrated and kept in a CO2-free incubator at 37°C. Meanwhile, bacteria were grown, and the macrophages were infected, as described above. After 6h post-infection, the media was removed, washed, and replaced with Seahorse XF media supplemented with glucose, pyruvate, and glutamine (Agilent) and stored in a CO2-free incubator at 37°C for one hour. For the mitochondrial stress test, oligomycin (1.5µM), carbonyl cyanide-4-(trifluoro ethoxy) phenylhydrazone (FCCP; 2µM), and Rotenone/ antimycin A (0.5µM) were prepared in Mito stress assay media and loaded into the appropriate Seahorse cartridge ports. The oxygen consumption rate (OCR) and extracellular acidification rate (ECAR) were measured every 3 min, and the appropriate compounds were injected sequentially at 18min intervals. ECAR and OCR were automatically calculated using the Wave software, and the graph was plotted in graph pad prism.

### Analysis of extracellular metabolites

Metabolite extraction for untargeted metabolomics was executed following the methodology published by Xu et al. (26). RAW264.7 cells were infected and incubated overnight with LPS, *S. anginosus* or *S. mitis*. After an overnight incubation, 1 ml of the cell culture media was collected and centrifuged at 5000 rpm for 10 min @ room temperature to pellet down bacteria and other debris. 100ul of the supernatant was added to the 900ul of metabolite extraction solution [Acetonitrile (75%), Methanol (25%), and Formic acid (0.2%)], which was then vortexed and centrifuged at 18000Xg for 10 min @ 4°C. Protein separation was performed on a Thermo Scientific Vanquish Duo UHPLC by The University of Arizona, Metabolomics Core. A Waters ACQUITY Premier HSS T3 column (1.8 μm, 2.1 mm x 150 mm) was used for reversed phase separation, and a Waters ACQUITY Premier BEH amide column (1.7 μm, 2.1 mm x 150 mm) was used for HILIC separation. For reversed phase seperation, the gradient was from 99% mobile phase A (0.1% formic acid in H_2_O) to 95% mobile phase B (0.1% formic acid in methanol) over 16 minutes. For HILIC separation, the gradient was the same as reversed phase with solvent A: 0.1% formic acid, 10 mM ammonium acetate, 90% acetonitrile, 10% H_2_O, and solvent B: 0.1% formic acid, 10 mM ammonium acetate, 50% acetonitrile, 50% H_2_O. Both columns were run at 45°C with a flow rate of 300 μL/min with an injection volume of 1 μL. Thermo Scientific Orbitrap Exploris 480 was used for data collection with a spray voltage of 3500 V for positive mode (reverse phase separation) and 2500 V for negative mode (HILIC separation) using the H-ESI source. The vaporizer temperature and ion transfer tube were both 350°C. Compounds were fragmented using data-dependent ms/ms with HCD collision energies of 20, 40, and 80%. Normalized peak areas were further analyzed using the online tool, Metaboanalyst 5.0.

### Statistical Analysis

GraphPad Prism software (version 9.4) was used for statistical analysis of the data and data visualization. One-way ANOVA followed by Dunnet’s post hoc analysis was carried out and all data are presented as mean ± SEM for biological triplicates.

## Results

### *S. anginosus* elicits significant morphological alterations indicating macrophage activation

RAW 264.7 macrophage cells were cultured in an antibiotic-free medium prior to infection with *S. anginosus* or *S. mitis* at different multiplicities of infection (MOI: 1, 10, and 50) for 6h and 24h, and macrophage growth morphology was observed using light microscopy after fixation (Supplementary figure. 2). Visualization of the fixed cells revealed significant changes in RAW264.7 cell morphology. Uninfected control cells exhibited rounded morphology, whereas 6h of LPS treatment resulted in elongated spindle-like morphology. Even greater elongated morphology was observed in a dose and time-dependent manner in *S. anginosus*-treated macrophages compared to LPS treated cells (Supplementary Figure. 2). In contrast, *S. mitis-*infected macrophages retained a rounded morphology at 6h and 24h of MOI 1 and MOI 10 treatment. However, at MOI 50, *S. mitis* induced slightly elongated morphology although this was not to the extent of *S. anginosus* infection at any measured MOI or time point (Supplementary Figure. 2). Furthermore, treatment of macrophages with LPS for 24h resulted in mixed, distorted, dead cell-like morphology. Thus, *S. anginosus* and *S. mitis* exhibit differential effects on macrophage morphology in a time and dose-dependent manner. Based on these morphometric findings, the MOI of 10 was chosen for further investigations. Light microscopy experiment was repeated with MOI 10 at 6h and followed by crystal violet staining for greater visualization of macrophage morphology. As shown in Figure. 1, time-dependent induction of pseudopodia-like structures was observed in macrophages treated with *S. anginosus* and *S. mitis* compared to LPS treated cells. Again, *S. anginosus* induced more prominent spindle-like structures in macrophages after 6h and 24h compared to other treatment groups. These observations suggest that S. *anginosus* activates macrophages more quickly and robustly than *S. mitis*, despite both being commensals of the oral cavity. Based on these morphometric findings, the MOI of 10 was chosen to study the innate immune response induced in RAW 264.7 macrophages by *S. anginosus* versus *S. mitis*.

**Figure 1:**
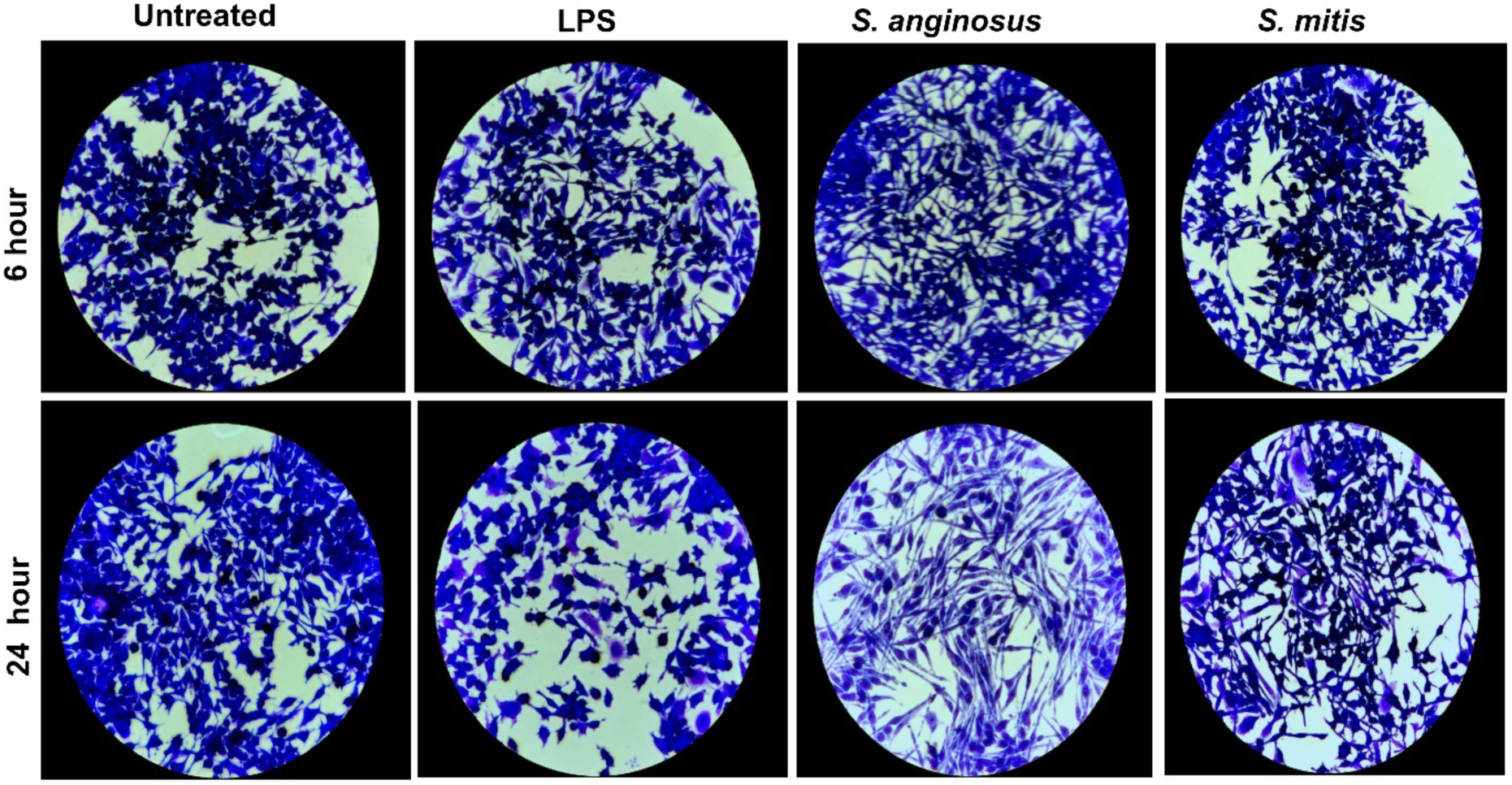
Crystal violet staining: Representative photograph of crystal violet staining of RAW264.7 macrophages (20X) depicts the macrophage phenotype that was analyzed after 6h (top) and 24h (bottom) post-infection at MOI 10. *S. anginosus*-infected macrophages showed a distinct phenotype with extended pseudopodia structures compared to untreated, LPS-treated, and *S. mitis*-infected macrophages.

### Giemsa staining reveals induction of binucleated giant macrophages by S. anginosus

Giemsa staining is a valuable tool for visualizing activated macrophages and assessing changes in cell size, as it provides contrast between cellular components, highlights morphological alterations, and facilitates quantification within macrophage-bacteria interaction studies. The present study utilized Giemsa staining to investigate the effect of *S. anginosus* and *S. mitis* on macrophage size as measured by cytoplasm-to-nuclei ratio. When activated, macrophages are large, 20-micron-diameter cells with an elongated nucleus containing a nucleolus, a nuclear-to-cytoplasmic ratio of less than one, and a much more complex cytoplasm (27). As shown in Figure. 2, *S. anginosus* infection resulted in distinct macrophage morphology associated with increased cellular size, more prominent cytoplasm, and the presence of binucleated nuclei (black arrow) located towards the cellular periphery (red arrow). These findings were observed in *S. anginosus* infected macrophages align with the morphological characteristics of activated macrophages, suggesting *S. anginosus* induces greater macrophage activation than *S. mitis*.

**Figure 2:**
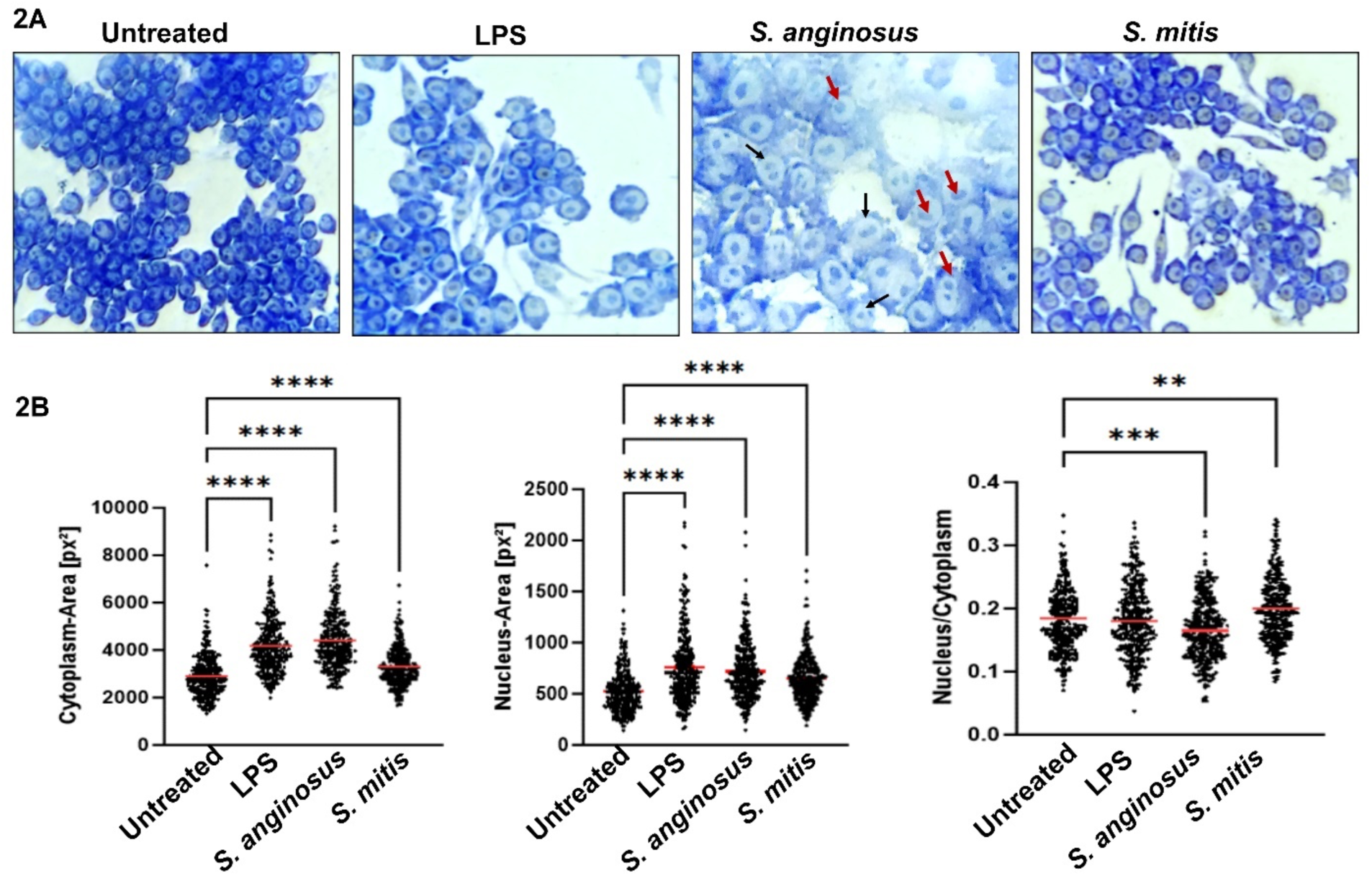
Giemsa Staining: (2A) Light microscopy images of RAW 264.7 (40X) incubated with LPS, *S. anginosus,* and *S. mitis* for 3h revealed that *S. anginosus* infection exhibited distinct macrophage phenotype with increased size like Giant cells, more prominent cytoplasm, and some of the nuclei are binucleated (black arrow). The peripheral position of nuclei (red arrow) is also observed in *S. anginosus* infected macrophages. (2B) Cell size quantification using NIS-Elements image analysis software. The nuclear to cytoplasmic ratio of activated macrophages is smaller due to expanded cytoplasm.

### *S. anginosus* elicits robust proinflammatory cytokine expression and production in RAW264.7 macrophages

Observing differentially altered cell morphology in macrophages exposed to *S. anginosus* and *S. mitis*, we proceeded to investigate its influence on inflammation by assessing the gene expression levels of the inflammatory cytokines TNF, IL6, and IL1β (Figure. 3A) following 6h post-infection and the protein levels of these cytokines in cell culture supernatants following 24h of infection by ELISA (Figure. 3B). *S. anginosus* treated macrophages displayed significantly elevated levels of proinflammatory cytokines compared to untreated macrophages, while *S. mitis* failed to elicit this robust immune response from macrophages.

**Figure 3:**
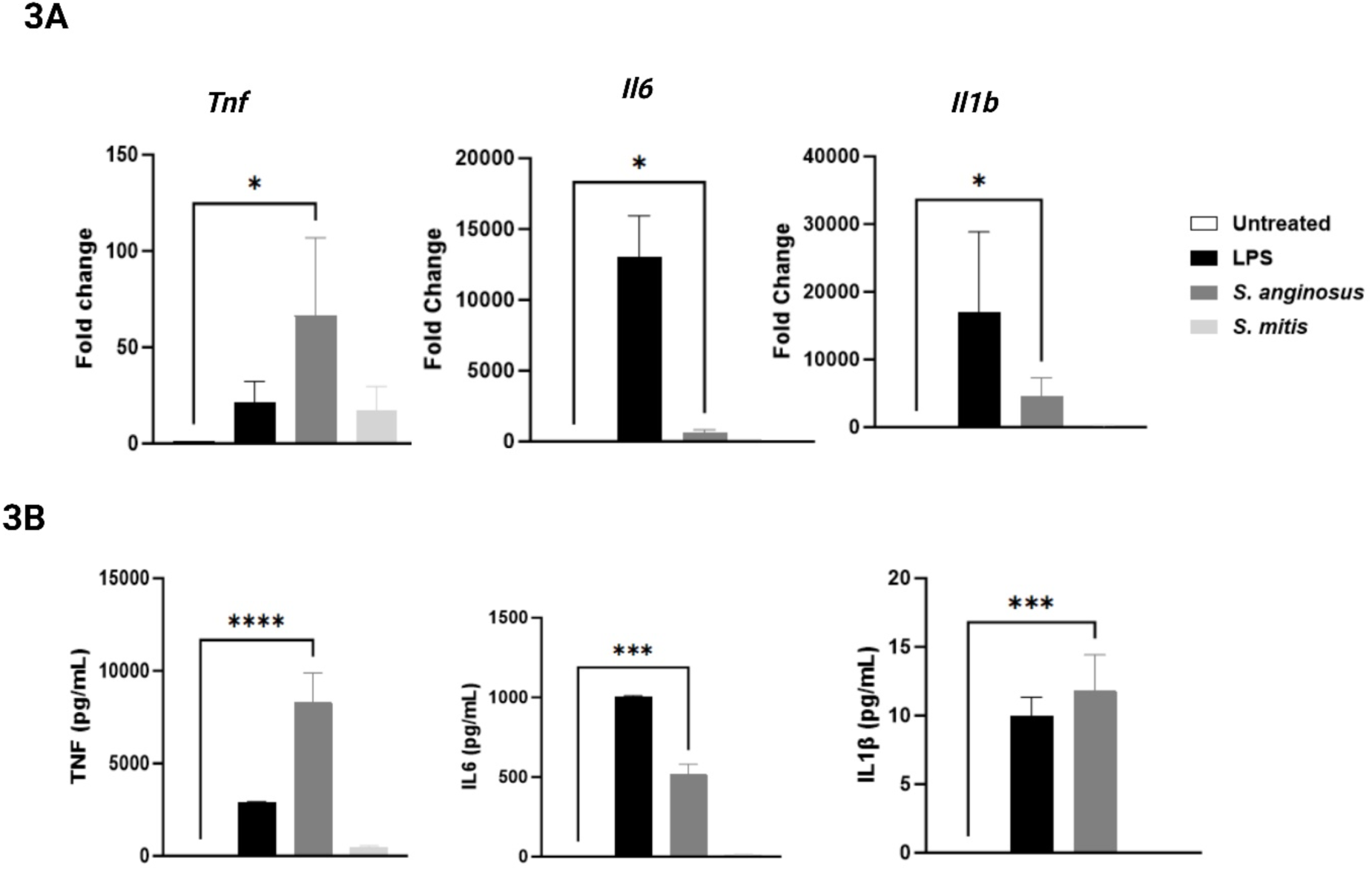
Proinflammatory cytokines expression. A) mRNA expression of cytokines from RAW 264.7 cells after 6h infection B) Cytokine secretion in cell culture supernatant as measured by ELISA after 24h post infection. The data are presented as Mean ± SEM. * P < 0.05 and ***P<0.0001 compared to untreated RAW 264.7 cells.

### *S. anginosus* drives macrophage activation through NF-κB stimulation

The NF-κB family of transcription factors are critical components of the immune response. NF-κB activation in stimulated cells is mediated by the phosphorylation and degradation of the inhibitory component IκBα, which allows for the release of p65. The released p65 is activated following phosphorylation, translating to the nucleus and transcriptional induction of several inflammatory cytokine genes, including TNF-α, IL-6, and IL-1β. To determine if *S. anginosus* and *S. mitis* also induce differential activation of NF-κB in macrophages, RAW267.4 macrophages were infected with *S. anginosus* or *S. mitis,* and the protein expression of total and phosphorylated p65 and IκBα were analyzed by western blot. As shown in Figure 4A, macrophages infected with *S. anginosus* for 1h resulted in significantly upregulated NF-κB activation, as evidenced by increased expression of phospho-p65 and phospho-IκBα compared to untreated controls. Moreover, no significant differences in NF-κB activation were observed between untreated macrophages and macrophages infected with *S. mitis* at this time point. These findings corroborated the increase in proinflammatory cytokines in macrophages infected *S. anginosus* compared to *S. mitis* (Figure 3A, B). The Nucleotide-binding domain and leucine-rich repeat-containing family, pyrin domain containing 3 (NLRP3), has gained attention as a vital facilitator of the innate immune system, playing a critical role in inflammatory responses during both sterile inflammatory insults and microbial infections, including several *Streptococcus* species (28,29). Compared to untreated control and *S. mitis* infected macrophages, *S. anginosus* infected macrophages significantly increased NLRP3 protein expression at 6h post-infection (Figure 4A, B). This pattern speculates a cascade wherein TLR4-activation primes NF-κB, subsequently stimulating NLRP3 and pro-IL-1β transcription.

**Figure 4:**
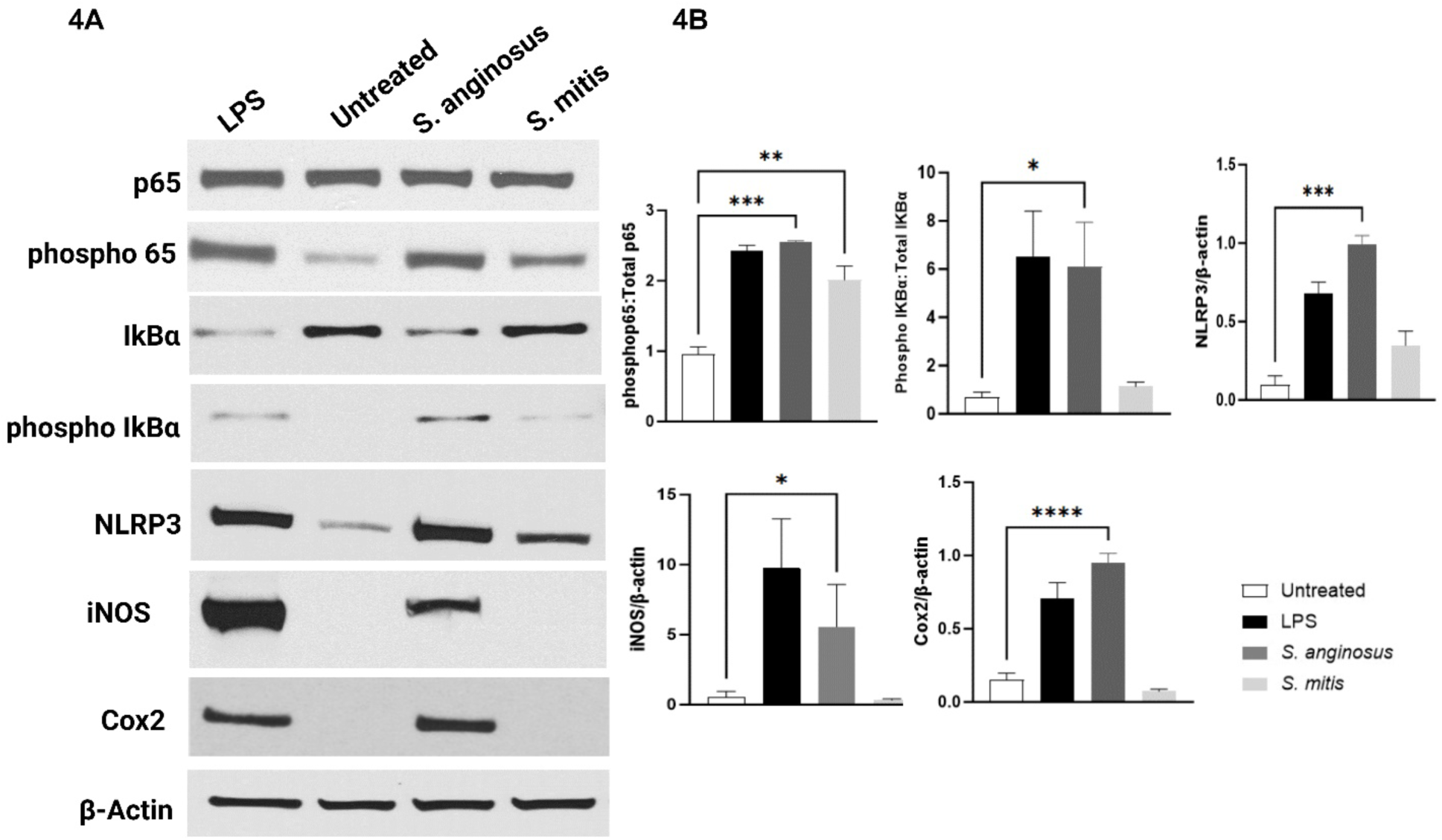
Protein expression of NF-κB and its downstream inflammatory mediators. 4A) Western blot analysis of RAW 264.7 cells treated with LPS, *S. anginosus* and *S. mitis* is shown. 4B). Densitometry analysis. The data are presented as Mean ± SEM. * P < 0.05 and ***P<0.0001 compared to untreated RAW 264.7 cells.

### *S. anginosus* induces inflammatory mediators including iNOS and Cox2

Inducible nitric oxide synthase (iNOS) and Cycloxygenase 2 (Cox2) are two inducible enzymes that generate inflammatory mediators, including nitric oxide and prostaglandins, respectively. The upregulation of iNOS and Cox2 during inflammation is regulated by the proinflammatory transcription factor NF-κB. Hence, we aimed to investigate if the expression of iNOS and Cox2 are differentially activated by *S. mitis* and *S. anginosus*. Accordingly, RAW 264.7 macrophages were exposed to *S. anginosus* or *S. mitis* infection for 6 hours, and the expression of iNOS and Cox2 was evaluated using qPCR and immunoblotting. Our results demonstrated that *S. anginosus* infection induced iNOS and Cox2 expression in macrophages, as depicted in Figure 4A and B. Conversely, *S. mitis* failed to induce the expression of these enzymes, which was consistent with the differential activation of NF-κB between these two strains of the genus *Streptococcus*. These observations demonstrate that when exposed to *S. anginosus*, macrophages mount a coordinated immune response against bacterial infection by increasing nitric oxide and prostaglandin production.

### S. anginosus-infected macrophages upregulated ACOD1, but not SDH

Inflammation triggers immune cell activation, shifting them from a quiescent state to an effector mode, marked by a significant revamp in cellular metabolism. This metabolic shift involves a profound restructuring of the citric acid cycle, a succession of enzymatic reactions within mitochondria. This cycle is the ultimate metabolic route for oxidizing multiple macronutrients such as carbohydrates, amino acids, and fats. Although the TCA cycle supports bioenergetics and biosynthesis, the role of the TCA cycle in macrophage polarization is not limited to ATP production. However, macrophages differentially utilize metabolic pathways to support maximal activation of phenotypes and effector functions (30). Classically activated M1 macrophages exhibit distinct modifications in the TCA cycle, characterized by interruptions after the metabolites citrate and succinate due to the upregulated expression of aconitate decarboxylase (ACOD1/IRG1) and inhibition of succinate dehydrogenase (SDH) enzymes. This metabolic reprogramming generates metabolites like itaconate and succinate, which can exert immunomodulatory effects by influencing gene expression, inflammatory responses, and immune regulation. After assessing NF-kB, iNOS, and Cox2, which are hallmarks of proinflammatory M1 macrophages, we next investigated the protein expression of ACOD1. This metabolic enzyme converts citrate to itaconate, and SDH, which catalyzes the conversion of succinate to fumarate. *S. anginosus* infection significantly upregulated ACOD1 protein expression in macrophages compared to untreated macrophages (Figure.5A). However, no significant alterations were observed in the expression of SDH in any of the treatment groups, suggesting a specific induction of ACOD1 in macrophages during *S. anginosus* infection. This observation signifies a potential role for TCA metabolic reprogramming in the context of *S. anginosus* induced macrophage activation.

**Figure 5:**
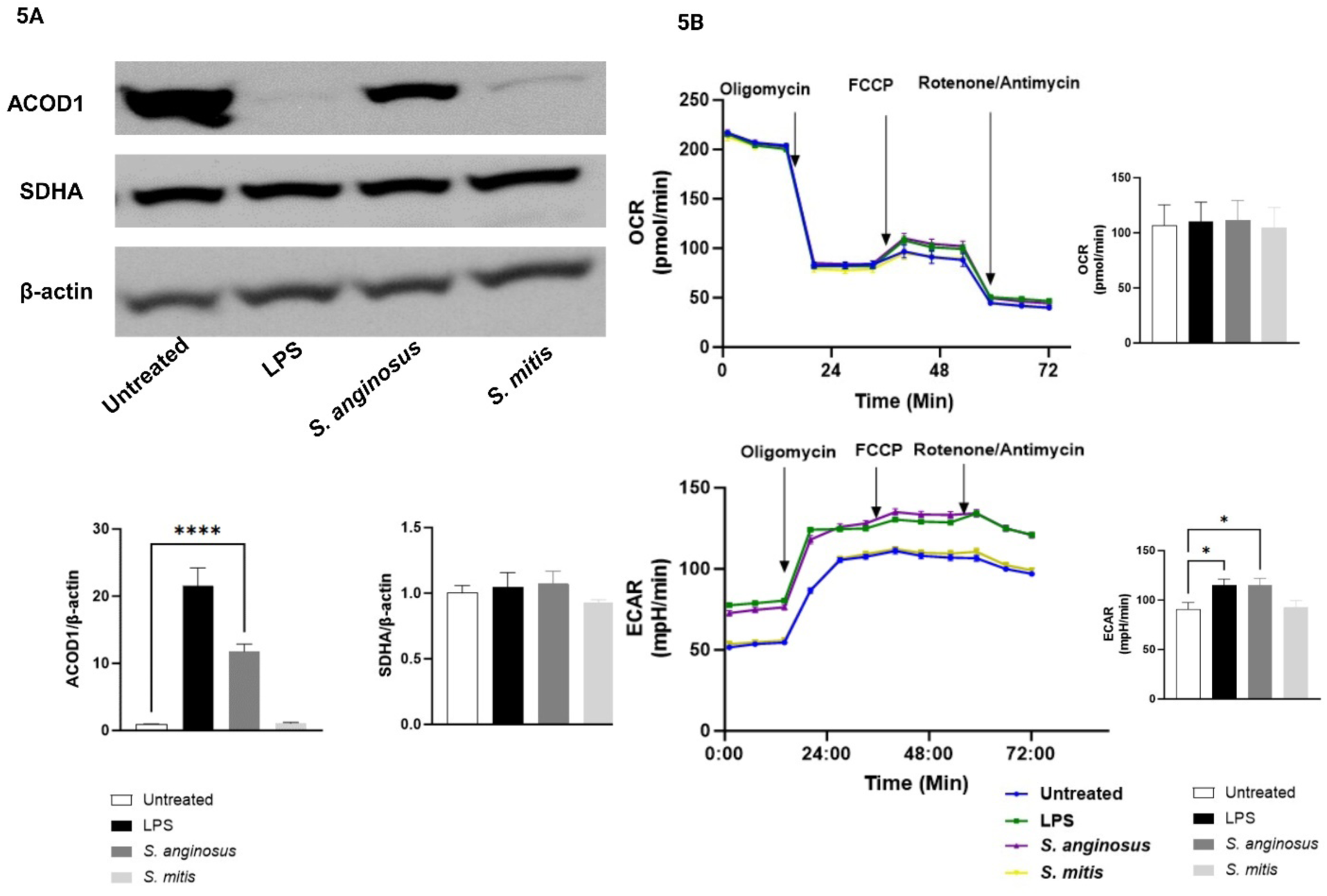
Metabolic analysis. 5A) Western blot and densitometry analysis of ACOD1 and SDHA. 5B) Seahorse xfe96 analysis of OCR and ECAR. The data are presented as Mean ± SEM. * P < 0.05 and ***P<0.0001 compared to untreated RAW 264.7 cells.

### S. anginosus, but not S. mitis infected macrophages showed increased ECAR

The measurement of extracellular rates of acidification and oxygen consumption rates provides a powerful way to assess the total energy metabolism of a cell. Oxygen consumption rate (OCR) quantifies how cells consume oxygen during mitochondrial respiration. In contrast, the extracellular acidification rate (ECAR) reflects the rate at which cells release protons (H+) into the extracellular environment, primarily due to the process of glycolysis (31). To assess the influence of *S. mitis* and *S. anginosus* on macrophage immunometabolism and bioenergetics, we evaluated alterations in OCR and ECAR after infecting the macrophages with these oral *Streptococcus* species for 6h. Our findings revealed that *S. anginosus* triggered heightened macrophage ECAR, akin to LPS treatment. Nevertheless, none of the experimental groups displayed noteworthy variations in OCR, implying that the 6h post-infection and LPS treatment had minimal impact on mitochondrial respiration and ATP production (Figure 5A). The increased ECAR, accompanied by a lack of alterations in OCR during *S. anginosus* infection, suggests a potential metabolic shift towards glycolysis induced by *S. anginosus,* which was absent in *S. mitis* infection (Figure 5B). This finding suggests *S. anginosus* and *S. mitis* have differential capacities to alter macrophage metabolism to drive cellular activation and proinflammatory signatures. Further investigation is required to understand this differential metabolic shift’s underlying mechanisms and functional consequences.

### *S. anginosus-*infected macrophages exhibit distinct immune-modulatory metabolite profiles and Itaconate-to-Succinate ratio

Many bacteria secrete small diffusible signal molecules to sense the local environmental conditions, including their population, and to synchronize multicellular behaviors. While several studies have focused on the intracellular concentration of various metabolites in RAW264.7 cells treated with LPS and other stimuli, we wanted to study the impact of microbial infection on extracellularly released metabolites, which may regulate microbial survival during host interactions. We performed untargeted metabolomics analysis on the supernatants of *S. anginosus* versus *S. mitis*-infected RAW264.7 macrophages and observed a demarcated cluster of metabolites in *S*. *anginosus* infected cells compared to *S. mitis*-infected or LPS-treated cells, as shown by the PCA plot (Figure.6A) and a unique metabolite profile as shown in the heat map (Figure.6B). The *S. anginosus* profile differences are highlighted by reduced glucose, arginine, and D-glutamine levels along with elevated lactic acid, amino acids, and other immune-related metabolites like acadesine, uric acid, D-aminoacids, and indole metabolites (Figure.6C).

**Figure 6:**
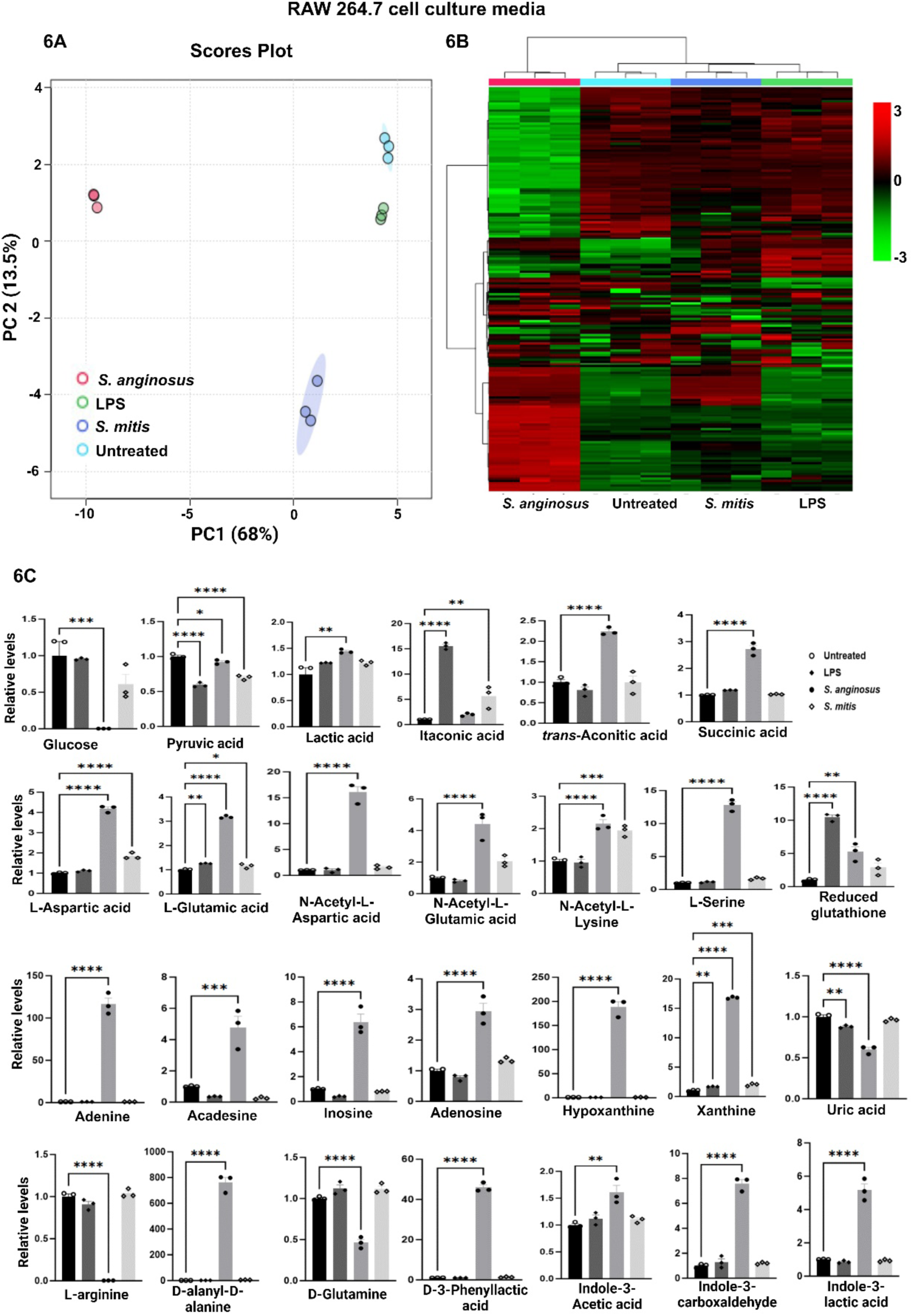
Extracellular metabolites analysis in *S. anginosus* and *S. mitis* infected RAW 264.7 cells: A) PCA Score Plot (n=3). B) Heat Map showing metabolite profile of experimental groups with *S. anginosus* displaying distinct metabolites profile. Color gradient: Red to black to green indicates relative metabolite level differences (scale adjacent). C) Bar Graph of critical metabolites related to infection and immunity, Analyzed in GraphPad Prism. Data presented as Mean ± SEM. * P < 0.05 and *** P < 0.0001 compared to untreated RAW 264.7 cells.

Notably, the upregulation of specific metabolites such as acetylated lysine and itaconate-to-succinate/transaconitic acid ratio modulation demonstrated distinctive immune responses induced by *S. anginosus* and *S. mitis*. Supplementary Table 1 shows the list of observed metabolites with their respective fold change in different experimental conditions. These findings provide insights into the unique immune-regulatory metabolite interactions triggered by these bacteria during infection, and the question of whether this ratio governs immune stimulation or suppression remains an uncharted domain, along with the potential role of trans-aconitic acid in this transition.

## Discussion

The intricate interplay between commensal microbiota and the mammalian immune system’s development and function maintains equilibrium. However, disruptions can contribute to disease, involving complex transitions from commensalism to pathogenicity influenced by various factors, including immune compromise and microbial traits, ultimately determining the nature and outcome of the host-microbe relationship (32). While host damage is often a requirement for the induction of a pathogen-specific immune response, the extent and nature of the damage incurred during the early stages and/or subsequent phases of interaction between the host and microbe determines the outcome of the host-microbe relationship (33). Oral streptococci or viridans group streptococci (VGS) are the first organisms to colonize oral surfaces, and most of them are typically deemed non-pathogenic due to their lack of virulence factors, like exotoxin production and due to their non-invasive potential found in other pathogenic organisms. Despite being traditionally considered non-pathogenic, they can also become opportunistic pathogens, evidenced by their involvement in a diverse range of clinical diseases, including infective endocarditis (34), bacteremia (35), and infections related to malignancy (36). A retrospective study conducted by Su et al. (37) identified the *S. anginosus* group (38.8%) and *S. mitis* group (22.8%) as the most common VGS species present in adult patients with a monomicrobial blood culture positive for VGS. Despite their clinical significance, understanding the nature of the immune response to these bacteria and the mechanisms underlying host defense remains unexplored. Understanding the host microbe interactions of these two *Streptococcus* species will help to understand infection dynamics host defense mechanisms, opening possibilities for treatment insights.

In this research, microscopic visualization highlighted *S. anginosus’* unique macrophage activation, evidenced by elongated cell structures in infected macrophages using crystal violet staining (38) and increased cytoplasmic area with peripheral nuclei through Giemsa staining (23), distinguishing it from *S. mitis*. Microbial pathogen-associated molecular patterns (PAMPs), such as gram-positive bacterial peptidoglycan, engage Toll-like receptors, frequently resulting in NF-kB activation; meanwhile, NLRP3 (NALP3, Cryopyrin, CIAS1, PYPAF1, CLR1.1), part of the cytosolic pattern-recognition receptor Nod-like receptor (NLR) family, recognizes microbial- and danger-associated molecular patterns, instigating innate immune responses. TLR-triggered NF-κB activation regulates NLRP3 transcription by binding to conserved NF-κB binding sites in the NLRP3 promoter, increasing NLRP3 expression (39). *S. anginosus* secretes an antigen termed SAA, a tyrosine tRNA synthetase (TyrRS) affiliated with the aminoacyl-tRNA synthetase family. The precise mechanism underlying SAA secretion remains unknown, as it lacks the conventional N-terminal signal peptide required for recognition by the secretion system (40). SAA has been demonstrated to dose-dependently stimulate NO production and induce the accumulation of NO synthetase mRNA in vitro in peritoneal exudate cells (PEC). SAA also induces the accumulation of TNF-α, IL-1β, and IL-6 mRNA (41). These reports correlate with the upregulation of NLRP3 and inflammatory cytokines, including TNFα, IL-6, and IL-1b, observed in *S. anginosus*-infected macrophages in the present study. Inducible nitric oxide synthase (iNOS) leads to the production of nitric oxide (NO), which acts as a signaling molecule to mediate diverse immune response and Cyclooxygenase 2 (Cox2), an inducible enzyme responds to inflammatory mediators by converting arachidonic acid to prostaglandins, thereby regulating inflammation in reaction to injury and infection. Both iNOs and Cox2 are associated with proinflammatory M1 macrophage with a potential to facilitate pathogen killing (42,43). During inflammation, the up-regulation of the inflammatory mediators, including iNOS and COX-2, is controlled by the proinflammatory transcription factor nuclear factor-κB (NF-kB). NF-κB represents a family of inducible transcription factors composed of five structurally related members, including NF-κB1 (also named p50), NF-κB2 (also named p52), RelA (also named p65), RelB, and c-Rel, which regulate an extensive array of genes involved in different processes of immunity and proinflammatory responses (44). This central player primarily mediates the coordination of the innate and adaptive immune response against invading microbial pathogens. *S. anginosus* significantly induced upregulation of phospho-p65 and phospho-IkBα and this upregulation is accompanied by the significantly increased levels of iNOS and Cox2 both at mRNA and protein levels in *S.anginosus* infected RAW264.7 cells as compared to untreated and *S. mitis*-infected macrophages, indicating potent activation of the canonical NF-κB pathway. However, *S. mitis*, which also secretes TyrRS, did not initiate a robust proinflammatory response in our experiments, which suggests that either the MOI or the time duration (or both) used in this study may need to be increased to observe a robust inflammatory response similar to *S. anginosus* infection.

Immunometabolism is crucial in host-pathogen interactions, with macrophage metabolism influencing infection outcomes. Immune cell disruption, which can occur during infection or tissue damage, releases signals from infected, dying, or stressed cells, inducing precise and timely inflammatory responses (45). To explore bacteria-macrophage metabolic interplay, we conducted non-targeted metabolomics on infected macrophage supernatants to identify secreted metabolites and real-time mito-stress tests for broader metabolic analysis and the overall metabolic profile of macrophages in response to *S. anginosus* and *S. mitis* is outlined in Figure 7. *S. anginosus* infected macrophages displayed a heightened consumption of glucose and increased production of lactic acid and ECAR. Upregulation of glucose utilization is a hallmark of classical macrophage activation, and increased glucose consumption fuels aerobic glycolysis, a consequence of diverting pyruvate (a product of glucose metabolism in the cytosol) towards lactate production rather than oxidation in the mitochondrial tricarboxylic acid (TCA) cycle (46). Following the detection of bacteria, macrophages switch their metabolism from oxidative respiration through the tricarboxylic acid cycle to high-rate aerobic glycolysis to meet the increased energy demands necessary to synthesize inflammatory cytokines (47,48). However, macrophage OCR and ATP production among the *Streptococcus*-infected and control macrophages did not differ significantly, suggesting that mitochondrial respiration is unaffected in any tested conditions at our experimental time point.

**Figure 7:**
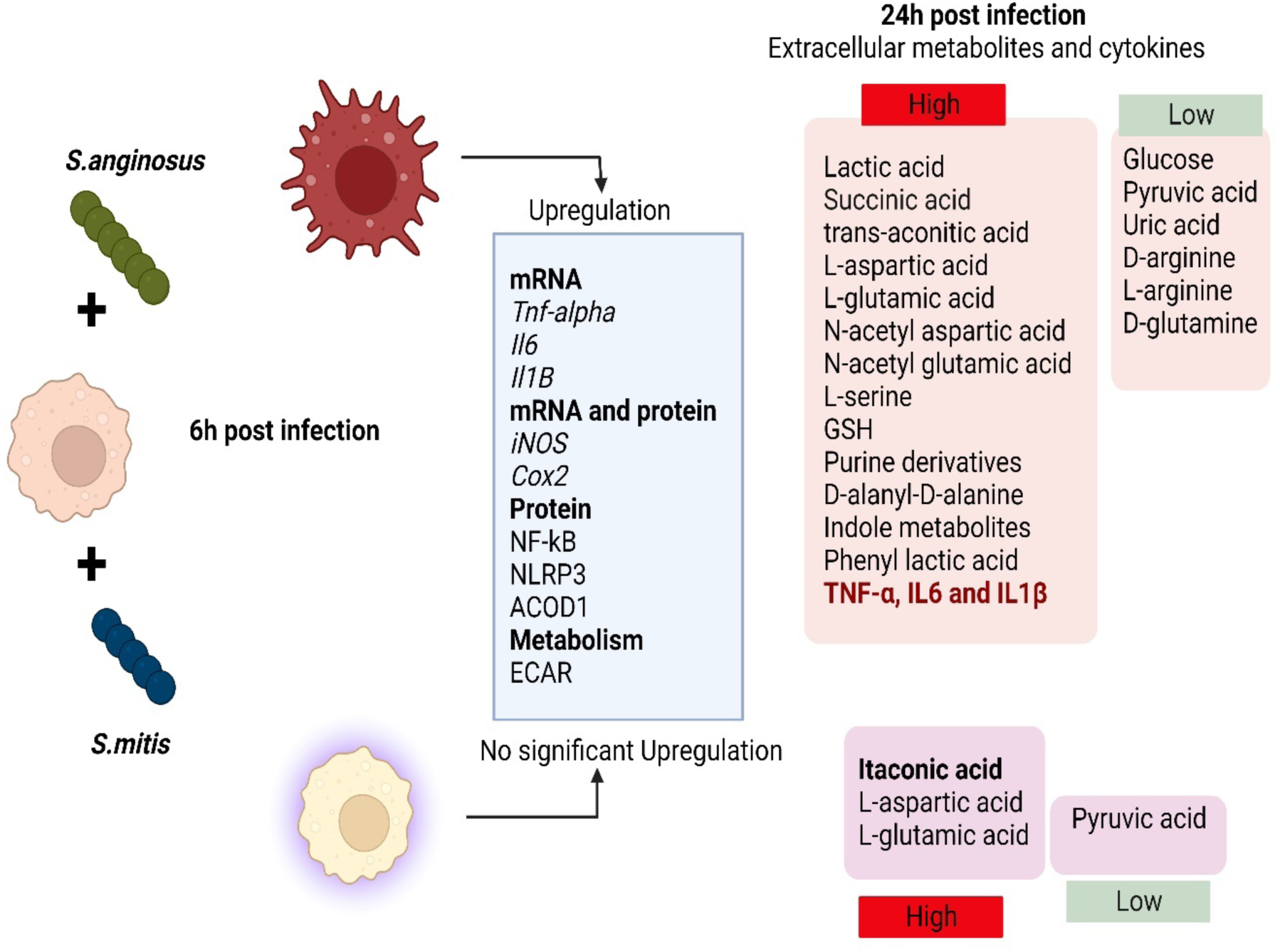
Graphical overview: Infection of macrophages by *S. anginosus* and *S. mitis* displays different morphological, immune and metabolic changes in 6h and 24h post infection.

In this study, *S. anginosus, S. mitis*, and LPS treatments significantly decreased macrophage extracellular pyruvate, a metabolite that enters diversified pathways in adaptation to cellular needs (49). *S. anginosus* exhibited a higher conversion of pyruvate to lactate in comparison to *S. mitis*. Alternatively, under other conditions, pyruvate might be channeled towards the TCA cycle as we found upregulation of itaconate in LPS and *S. mitis* treated macrophage media but not from *S.anginosus* 24h post-infection. This redirection could potentially enhance NADH generation, fueling the oxidative phosphorylation observed in resting cells. Glucose levels remained stable in LPS- and *S. mitis*-treated macrophages, consistent with the findings of Newsholme et al., who reported a low rate of glucose utilization by ‘resting’ macrophages in vitro, with a sharp increase in glycolysis during phagocytosis or heightened secretory activity. Sun et al. (50) demonstrated that the impact of particulate matter on mitochondrial oxygen consumption varies with time, suggesting that exploring metabolic flux over more extended incubation periods could modify the OCR. This avenue is worth further investigation, especially regarding *S. mitis* versus *S. anginosus* infection.

The Warburg effect, marked by impaired mitochondrial respiration, is typically linked to M1 proinflammatory macrophages, often accompanied by a disrupted mitochondrial Citric acid cycle (TCA cycle). It is proposed that LPS stimulation downregulates the mRNA levels of the enzyme isocitrate dehydrogenase (IDH), creating a cellular pool of citrate that supports fatty acid synthesis and the generation of lipid intermediates like prostaglandins that promote inflammation. Decreases in IDH activity may also contribute to the accumulation of citrate-related metabolites, supporting the production of itaconate (51), a metabolic byproduct derived from cis-aconitic acid in the TCA cycle, which is catalyzed by cis-aconitate decarboxylase (IRG1/CAD/ACOD1). In proinflammatory cells, itaconate serves two critical functions. First, it inhibits bacterial isocitrate lyase (52), disrupting the glyoxylate shunt, a critical metabolic pathway that impairs bacterial growth in carbohydrate-limiting conditions. Secondly, it inhibits succinate dehydrogenase (SDH), an enzyme that converts succinate to fumarate within the host cell, leading to succinate accumulation (53). *S. anginosus* infection exhibited temporal dynamics on ACOD1 expression (upregulation following 6h post-infection) and Itaconate production (no significant increase in 24h post-infection). Further, *S.anginosus* infection also causes a surge in the TCA cycle metabolite succinate. While succinate and itaconate accumulation are hallmarks of proinflammatory macrophages, we found elevated extracellular itaconate in LPS-stimulated and *S. mitis-*infected macrophages at 24h, which differed from *S. anginosus* infection. Furthermore, while LPS- and IFN-γ-driven macrophage activation increases itaconate levels, which aligns with our observations regarding LPS, *S. anginosus* infection induces distinct metabolic changes to trigger inflammatory responses (54, 55).

Succinate is an important metabolite in both host and microbial processes. Although normally regarded as an intermediate, succinate is observed to accumulate in certain pathophysiological situations, especially in areas of inflammation and metabolic stress. Succinate is found in diverse tissues and immune cells and contributes to proinflammatory responses and stress-induced immune activation. Microbial disruptions in gut microbiota metabolism and cross-feeding can lead to succinate accumulation, especially with dietary changes, indigestible carbohydrate consumption, and antibiotic use, resulting in elevated fecal succinate levels in the elderly and various animal models as reviewed by Connors et al. (57). In combination with LPS stimulation, succinate enhances HIF-1α stability, driving IL-1β transcription (56). When released extracellularly, succinate signals through upregulated SUCNR1/GPR91 and magnifies IL-1β production, establishing a macrophage activation loop (58). Conversely, studies using SUCNR1-deficient peritoneal macrophages revealed similar LPS-induced cytokine profiles as wild-type controls (59). Moreover, Keiran et al. (60) demonstrated that succinate/SUCNR1 signaling promotes an anti-inflammatory macrophage phenotype, emphasizing succinate’s dual role as a proinflammatory danger signal and stress-induced alarmin alerting the immune system. *Staphylococcus aureus*, another Gram-positive bacterium, is a weak inducer of IL-1β due to its ability to promote higher itaconate and lower succinate levels (61), resembling the effect of *S. mitis* in the current study. The source and impact of elevated extracellular succinate in macrophages in response to *S. anginosus* infection warrants further investigation.

In this study, supernatants from *S. anginosus*-infected macrophages displayed markedly elevated levels of trans-aconitic acid(tAA), an isomer of cis-aconitate and a precursor for itaconate. tAA is found in *Pseudomonas* (62) and certain plants (63). *Bacillus thuringiensis-*produced tAA exhibits antibacterial effects against *Meloidogyne incognita* (64). tAA is reported to be a potent aconitase (ACO2) inhibitor, an enzyme that converts citrate to isocitrate via cis-aconitate. Disabling ACO2 activity or expression can be employed to truncate citrate flux, permitting its extrusion outside the mitochondria for lipogenesis, which plays a vital role in carcinogenesis. Moreover, the loss of mitochondrial ACO2 promotes colorectal cancer progression via lipid remodeling (65). In the presence of high NO concentrations in LPS+IFNγ-stimulated macrophages, carbon entry into the TCA cycle via PDH is inhibited, and ACO2 is suppressed. Enhanced compensatory carboxylation and glutaminolysis lead to citrate accumulation and itaconate limitation (66). In this context, in addition to *S. anginosus* secretion of SAA/TyrS, upregulated tAA in *S.anginosus* infection could also interfere with ACO2, which would be in line with the observed limited itaconate secretion and reduced glutamine at 24-hour post-infection.

L-arginine (L-arg) is a versatile amino acid and a central intestinal metabolite in mammalian and microbial organisms and serves as a substrate for intestinal and microbial cells that colonize the largest interface at which our body crosstalks to its microbiota. L-arg deprivation prolongs bacterial persistence and perpetuates chronic inflammation (67). D-Glutamine, an enantiomer of the conditionally essential amino acid L-glutamine, is reduced in the serum of rats in a model of acute pancreatitis (68) and in patients with hepatocellular carcinoma (69). *S. anginosus-*infected macrophages displayed significantly decreased levels of D-glutamine and L arginine, suggesting S. *anginosus* has a greater capacity to utilize these amino acids compared to *S. mitis*, which may contribute to S. *anginosus* pathogenicity. These novel findings warrant further investigation to better understand how *S. anginosus* mediates immune responses in the context of oral biology and squamous cell carcinoma progression.

In summary, here we found that *S. anginosus* triggers a proinflammatory response similar to LPS stimulation at 6 hours post-infection yet induces a distinct extracellular metabolite profile compared to LPS alone. Conversely, *S. mitis* induces a diminished inflammatory response compared to LPS and *S. anginosus* infection but promotes the release of immune-regulatory metabolites similar to LPS treatment. These distinct inflammatory and metabolic shifts in macrophages during *S. anginosus* and *S. mitis* infection warrant further investigation. Considering the heightened extracellular levels of trans-aconitic acid and succinic acid induced during *S. anginosus* infection, *S. anginosus* may influence macrophage immunometabolism through engagement of the tAA-ACO2-NO axis. This warrants further exploration using primary cells to evaluate the impact of *S. anginosus* on these metabolic pathways during inflammatory signaling and immune activation.

## ACKNOWLEDGEMENTS

This work was supported by the University of Arizona Health Sciences Strategic Initiatives - Personalized Defense sanctioned under the Immune-Microbe Interface in Health and Disease (IMIHD) group. We greatly acknowledge the Analytical & Biological Mass Spectrometry Facility at The University of Arizona, Tucson, AZ, for metabolite analysis. We sincerely thank Dr. Rick G. Schnellmann, Dean, College of Pharmacy, The University of Arizona, for his timely support with seahorse analysis. Additionally, we would like to thank Merchant Lab, University of Arizona Cancer Center, for generously providing access to their microscopy facility. Illustrations were created with Biorender.com.

The Authors declare no conflicts of interest.

## References

1. Deo PN, Deshmukh R. 2019. Oral microbiome: Unveiling the fundamentals. J Oral Maxillofac Pathol 23:122–128. 10.4103/jomfp.JOMFP_304_18.

2. Maeda Y, Goldsmith CE, Coulter WA, Mason C, Dooley J, Lowery C, Moore JE. 2010. The viridans group streptococci. Rev Med Microbiol 21:69–79. 10.1097/MRM.0b013e32833c68fa

3. Ammann TW, Belibasakis GN, Thurnheer T. 2013. Impact of early colonizers on in vitro subgingival biofilm formation. PloS one 8:e83090. 10.1371/journal.pone.0083090

4. Abranches J, Zeng L, Kajfasz JK, Palmer SR, Chakraborty B, Wen ZT, Richards VP, Brady LJ, Lemos JA. 2018. Biology of Oral Streptococci. Microbiol Spectr 6: 10.1128/microbiolspec.GPP3-0042-2018. https://doi.org/10.1128/microbiolspec.GPP3-0042-2018

5. Schmidt BL, Kuczynski J, Bhattacharya A, Huey B, Corby PM, Queiroz EL, Nightingale K, Kerr AR, DeLacure MD, Veeramachaneni R, Olshen AB, Albertson DG. 2014. Changes in abundance of oral microbiota associated with oral cancer. PLoS One 9:e98741. 10.1371/journal.pone.0098741

6. Al Majid F, Aldrees A, Barry M, Binkhamis K, Allam A, Almohaya A. (2020). Streptococcus anginosus group infections: Management and outcome at a tertiary care hospital. J Infect Public Health 13:1749–1754. 10.1016/j.jiph.2020.07.017

7. Pilarczyk-Zurek M, Sitkiewicz I, Koziel J. 2022. The Clinical View on Streptococcus anginosus Group – Opportunistic Pathogens Coming Out of Hiding. Front Microbiol 13:956677. 10.3389/fmicb.2022.956677.

8. Morita E, Narikiyo M, Yano A, Nishimura E, Igaki H, Sasaki H, Terada M, Hanada N, Kawabe R. 2003. Different frequencies of Streptococcus anginosus infection in oral cancer and esophageal cancer. Cancer Sci 94:492–496. 10.1111/j.1349-7006.2003.tb01471.x

9. Masood U, Sharma A, Lowe D, Khan R, Manocha D. 2016. Colorectal Cancer Associated with Streptococcus anginosus Bacteremia and Liver Abscesses. Case Rep Gastroenterol 10:769–774. 10.1159/000452757

10. Tateda M, Shiga K, Saijo S, Sone M, Hori T, Yokoyama J, Matsuura K, Takasaka T, Miyagi T. 2000. Streptococcus anginosus in head and neck squamous cell carcinoma: implication in carcinogenesis. Int J Mol Med 6:699–703. 10.3892/ijmm.6.6.699

11. Xu Y, Jia Y, Chen L, Gao J, Yang D. 2021. Effect of Streptococcus anginosus on biological response of tongue squamous cell carcinoma cells. BMC Oral Health 21:141. 10.1186/s12903-021-01505-3.

12. Mager DL, Haffajee AD, Devlin PM, Norris CM, Posner MR, Goodson JM. 2005. The salivary microbiota as a diagnostic indicator of oral cancer: a descriptive, non-randomized study of cancer-free and oral squamous cell carcinoma subjects. J Transl Med 3:27. 10.1186/1479-5876-3-27.

13. Sharma AV, Masood U, Kahlon AS, Pattar S, Iqbal S, Lehmann D. 2016. *Streptococcus mitis* Bacteremia and Endocarditis: An Early Sign in Pre-Cancerous Colon Polyps: 1366. Am J Gastroenterol 111: S616. https://api.semanticscholar.org/CorpusID:208443761

14. Narikiyo M, Tanabe C, Yamada Y, Igaki H, Tachimori Y, Kato H, Muto M, Montesano R, Sakamoto H, Nakajima Y, Sasaki H. 2004. Frequent and preferential infection of Treponema denticola, Streptococcus mitis, and Streptococcus anginosus in esophageal cancers. Cancer Sci 95:569–574. 10.1111/j.1349-7006.2004.tb02488.x

15. Tsai MS, Chen YY, Chen WC, Chen MF. 2022. Streptococcus mutans promotes tumor progression in oral squamous cell carcinoma. J Cancer 13:3358–3367. 10.7150/jca.73310.

16. Lim JJ, Grinstein S, Roth Z. 2017. Diversity and Versatility of Phagocytosis: Roles in Innate Immunity, Tissue Remodeling, and Homeostasis. Front Cell Infect Microbiol 7:191. 10.3389/fcimb.2017.00191.

17. Yunna C, Mengru H, Lei W, Weidong C. 2020. Macrophage M1/M2 polarization. Eur J Pharmacol 877:173090. 10.1016/j.ejphar.2020.173090

18. Benoit M, Desnues B, Mege JL. 2008. Macrophage polarization in bacterial infections. J Immunol 181:3733–3739. 10.4049/jimmunol.181.6.3733.

19. Menghani SV, Rivera A, Neubert M, Hagerty JR, Lewis L, Galgiani JN, Jolly ER, Alvin JW, Johnson MDL. 2021. Demonstration of N,N-Dimethyldithiocarbamate as a Copper-Dependent Antibiotic against Multiple Upper Respiratory Tract Pathogens. Microbiol Spectr 9:e0077821. 10.1128/Spectrum.00778-21

20. Martis PC, Dudley AT, Bemrose MA, Gazda HL, Smith BH, Gazda LS. 2018. MEF2 plays a significant role in the tumor inhibitory mechanism of encapsulated RENCA cells via EGF receptor signaling in target tumor cells. BMC Cancer 18:1217. 10.1186/s12885-018-5128-5.

21. Zinser GM, McEleney K, Welsh J. 2003. Characterization of mammary tumor cell lines from wild type and vitamin D3 receptor knockout mice. Mol Cell Endocrinol 200:67–80. 10.1016/s0303-7207(02)00416-1.

22. Feoktistova M, Geserick P, Leverkus M. 2016. Crystal Violet Assay for Determining Viability of Cultured Cells. Cold Spring Harb Protoc 2016:pdb.prot087379. 10.1101/pdb.prot087379.

23. Nicola AM, Casadevall A. 2012. In vitro measurement of phagocytosis and killing of Cryptococcus neoformans by macrophages. Methods Mol Biol 844:189–197. https://api.semanticscholar.org/CorpusID:37069037.

24. Toni LS, Garcia AM, Jeffrey DA, Jiang X, Stauffer BL, Miyamoto SD, Sucharov CC. 2018. Optimization of phenol-chloroform RNA extraction. MethodsX 5:599–608. https://api.semanticscholar.org/CorpusID:37069037.

25. Wilson JE, Petrucelli AS, Chen L, Koblansky AA, Truax AD, Oyama Y, Rogers AB, Brickey WJ, Wang Y, Schneider M, Mühlbauer M, Chou WC, Barker BR, Jobin C, Allbritton NL, Ramsden DA, Davis BK, Ting JP. 2015. Inflammasome-independent role of AIM2 in suppressing colon tumorigenesis via DNA-PK and Akt. Nat Med 21:906–13. 10.1038/nm.3908

26. Xu Y, Jia Y, Chen L, Gao J, Yang D. 2021. Effect of Streptococcus anginosus on biological response of tongue squamous cell carcinoma cells. BMC Oral Health 21:141. 10.1186/s12903-021-01505-3.

27. Anderson KM, Ondrey FG, Harris JE. 1989. 5,8,11,14-eicosatetraynoic acid-induced differentiation of U937 cells. Ann Clin Lab Sci 19: 92–100.

28. Harder J, Franchi L, Muñoz-Planillo R, Park JH, Reimer T, Núñez G. 2009. Activation of the Nlrp3 inflammasome by Streptococcus pyogenes requires streptolysin O and NF-kappa B activation but proceeds independently of TLR signaling and P2X7 receptor. J Immunol 183:5823–5829. 10.4049/jimmunol.0900444.

29. Witzenrath M, Pache F, Lorenz D, Koppe U, Gutbier B, Tabeling C, Reppe K, Meixenberger K, Dorhoi A, Ma J, Holmes A, Trendelenburg G, Heimesaat MM, Bereswill S, van der Linden M, Tschopp J, Mitchell TJ, Suttorp N, Opitz B. 2011. The NLRP3 inflammasome is differentially activated by pneumolysin variants and contributes to host defense in pneumococcal pneumonia. J Immunol 187:434–440. 10.4049/jimmunol.1003143

30. Diskin C, Pålsson-McDermott EM. 2018. Metabolic Modulation in Macrophage Effector Function. Front Immunol 9:270. 10.3389/fimmu.2018.00270.

31. Mookerjee SA, Gerencser AA, Nicholls DG, Brand MD. 2017. Quantifying intracellular rates of glycolytic and oxidative ATP production and consumption using extracellular flux measurements. J Biol Chem 292:7189–7207. 10.1074/jbc.M116.774471.

32. Thakur A, Mikkelsen H, Jungersen G. 2019. Intracellular Pathogens: Host Immunity and Microbial Persistence Strategies. J Immunol Res 2019:1356540. 10.1155/2019/1356540

33. Casadevall A, Pirofski LA. 2000. Host-pathogen interactions: basic concepts of microbial commensalism, colonization, infection, and disease. Infect Immun 68:6511–6518. 10.1128/IAI.68.12.6511-6518.2000.

34. Kim SL, Gordon SM, Shrestha NK. 2018. Distribution of streptococcal groups causing infective endocarditis: a descriptive study. Diagn Microbiol Infect Dis 91:269–272. 10.1016/j.diagmicrobio.2018.02.015.

35. Tran MP, Caldwell-McMillan M, Khalife W, Young VB. 2008. Streptococcus intermedius causing infective endocarditis and abscesses: a report of three cases and review of the literature. BMC Infect Dis 8:154. 10.1186/1471-2334-8-154.

36. Yu L, Maishi N, Akahori E, Hasebe A, Takeda R, Matsuda AY, Hida Y, Nam JM, Onodera Y, Kitagawa Y, Hida K. 2022. The oral bacterium Streptococcus mutans promotes tumor metastasis by inducing vascular inflammation. Cancer Sci 113:3980–3994. 10.1111/cas.15538.

37. Su TY, Lee MH, Huang CT, Liu TP, Lu JJ. 2018. The clinical impact of patients with bloodstream infection with different groups of Viridans group streptococci by using matrix-assisted laser desorption ionization-time of flight mass spectrometry (MALDI-TOF MS). Medicine (Baltimore) 97:e13607. 10.1097/MD.0000000000013607

38. Liu HF, Zhang HJ, Hu QX, Liu XY, Wang ZQ, Fan JY, Zhan M, Chen FL. 2012. Altered polarization, morphology, and impaired innate immunity germane to resident peritoneal macrophages in mice with long-term type 2 diabetes. J Biomed Biotechnol 2012:867023. 10.1155/2012/867023.

39. Qiao Y, Wang P, Qi J, Zhang L, Gao C. 2012. TLR-induced NF-κB activation regulates NLRP3 expression in murine macrophages. FEBS Lett 586:1022–1026. 10.1016/j.febslet.2012.02.045

40. Shimoyama Y, Ishikawa T, Kodama Y, Kimura S, Sasaki M. 2020. Tyrosine tRNA synthetase as a novel extracellular immunomodulatory protein in Streptococcus anginosus. FEMS Microbiol Lett 367:fnaa153. DOI: 10.1093/femsle/fnaa153. PMID: 32926111.

41. Sasaki M, Ohara-Nemoto Y, Tajika S, Kobayashi M, Yamaura C, Kimura S. 2001. Antigenic characterisation of a novel Streptococcus anginosus antigen that induces nitric oxide synthesis by murine peritoneal exudate cells. J Med Microbiol 50: 952–958. 10.1099/0022-1317-50-11-952.

42. Orecchioni M, Ghosheh Y, Pramod AB, Ley K. 2019. Macrophage Polarization: Different Gene Signatures in M1(LPS+) vs. Classically and M2(LPS-) vs. Alternatively Activated Macrophages. Front Immunol 10:1084. 10.3389/fimmu.2019.01084

43. Ren H, Chen X, Jiang F, Li G. 2020. Cyclooxygenase-2 Inhibition Reduces Autophagy of Macrophages Enhancing Extraintestinal Pathogenic Escherichia coli Infection. Front Microbiol 11:708. 10.3389/fmicb.2020.00708.

44. Oeckinghaus A, Ghosh S. 2009. The NF-kappaB family of transcription factors and its regulation. Cold Spring Harb Perspect Biol 1:a000034. 10.1101/cshperspect.a000034.

45. Gaber T, Strehl C, Buttgereit F. 2017. Metabolic regulation of inflammation. Nat Rev Rheumatol 13:267–279. 10.1038/nrrheum.2017.37.

46. Jung J, Zeng H, Horng T. 2019. Metabolism as a guiding force for immunity. Nat Cell Biol 21:85–93. 10.1038/s41556-018-0217-x.

47. He W, Heinz A, Jahn D, Hiller K. 2021. Complexity of macrophage metabolism in infection. Curr Opin Biotechnol 68:231–239. 10.1016/j.copbio.2021.01.020

48. Rosenberg G, Riquelme S, Prince A, Avraham R. 2022. Immunometabolic crosstalk during bacterial infection. Nat Microbiol 7:497–507. 10.1038/s41564-022-01080-5.

49. Prochownik EV, Wang H. 2021. The Metabolic Fates of Pyruvate in Normal and Neoplastic Cells. Cells 10:762. 10.3390/cells10040762

50. Sun KA, Li Y, Meliton AY, Woods PS, Kimmig LM, Cetin-Atalay R, Hamanaka RB, Mutlu GM. 2020.Endogenous itaconate is not required for particulate matter-induced NRF2 expression or inflammatory response. Elife 9:e54877. 10.7554/eLife.54877.

51. Jha AK, Huang SC, Sergushichev A, Lampropoulou V, Ivanova Y, Loginicheva E, Chmielewski K, Stewart KM, Ashall J, Everts B, Pearce EJ, Driggers EM, Artyomov MN. 2015. Network integration of parallel metabolic and transcriptional data reveals metabolic modules that regulate macrophage polarization. Immunity 42:419–30. 10.1016/j.immuni.2015.02.005

52. McFadden BA, Purohit S. 1977. Itaconate, an isocitrate lyase-directed inhibitor in Pseudomonas indigofera. J Bacteriol 131:136–44. 10.1128/jb.131.1.136-144.1977.

53. Cordes T, Wallace M, Michelucci A, Divakaruni AS, Sapcariu SC, Sousa C, Koseki H, Cabrales P, Murphy AN, Hiller K, Metallo CM. 2016. Immunoresponsive Gene 1 and Itaconate Inhibit Succinate Dehydrogenase to Modulate Intracellular Succinate Levels. J Biol Chem 291:14274–14284. 10.1074/jbc.M115.685792.

54. Strelko CL, Lu W, Dufort FJ, Seyfried TN, Chiles TC, Rabinowitz JD, Roberts MF. 2011. Itaconic acid is a mammalian metabolite induced during macrophage activation. J Am Chem Soc 133:16386–16389. 10.1021/ja2070889

55. Lampropoulou V, Sergushichev A, Bambouskova M, Nair S, Vincent EE, Loginicheva E, Cervantes-Barragan L, Ma X, Huang SC, Griss T, Weinheimer CJ, Khader S, Randolph GJ, Pearce EJ, Jones RG, Diwan A, Diamond MS, Artyomov MN. 2016. Itaconate Links Inhibition of Succinate Dehydrogenase with Macrophage Metabolic Remodeling and Regulation of Inflammation. Cell Metab 24:158–166. 10.1016/j.cmet.2016.06.004.

56. Tannahill GM, Curtis AM, Adamik J, Palsson-McDermott EM, McGettrick AF, Goel G, Frezza C, Bernard NJ, Kelly B, Foley NH, Zheng L, Gardet A, Tong Z, Jany SS, Corr SC, Haneklaus M, Caffrey BE, Pierce K, Walmsley S, Beasley FC, Cummins E, Nizet V, Whyte M, Taylor CT, Lin H, Masters SL, Gottlieb E, Kelly VP, Clish C, Auron PE, Xavier RJ, O’Neill LA. 2013. Succinate is an inflammatory signal that induces IL-1β through HIF-1α. Nature 496:238–242. 10.1038/nature11986

57. Connors J, Dawe N, Van Limbergen J. 2018. The Role of Succinate in the Regulation of Intestinal Inflammation. Nutrients.11:25. 10.3390/nu11010025

58. Littlewood-Evans A, Sarret S, Apfel V, Loesle P, Dawson J, Zhang J, Muller A, Tigani B, Kneuer R, Patel S, Valeaux S, Gommermann N, Rubic-Schneider T, Junt T, Carballido JM. 2016. GPR91 senses extracellular succinate released from inflammatory macrophages and exacerbates rheumatoid arthritis. J Exp Med 213:1655–1662. 10.1084/jem.20160061.

59. van Diepen JA, Robben JH, Hooiveld GJ, Carmone C, Alsady M, Boutens L, Bekkenkamp-Grovenstein M, Hijmans A, Engelke UFH, Wevers RA, Netea MG, Tack CJ, Stienstra R, Deen PMT. 2017. SUCNR1-mediated chemotaxis of macrophages aggravates obesity-induced inflammation and diabetes. Diabetologia 60:1304–1313. 10.1007/s00125-017-4261-z.

60. Keiran N, Ceperuelo-Mallafré V, Calvo E, Hernández-Alvarez MI, Ejarque M, Núñez-Roa C, Horrillo D, Maymó-Masip E, Rodríguez MM, Fradera R, de la Rosa JV, Jorba R, Megia A, Zorzano A, Medina-Gómez G, Serena C, Castrillo A, Vendrell J, Fernández-Veledo S. 2019. SUCNR1 controls an anti-inflammatory program in macrophages to regulate the metabolic response to obesity. Nat Immunol 20:581–592. 10.1038/s41590-019-0372-7.

61. Tomlinson KL, Lung TWF, Dach F, Annavajhala MK, Gabryszewski SJ, Groves RA, Drikic M, Francoeur NJ, Sridhar SH, Smith ML, Khanal S, Britto CJ, Sebra R, Lewis I, Uhlemann AC, Kahl BC, Prince AS, Riquelme SA. 2021. Staphylococcus aureus induces an itaconate-dominated immunometabolic response that drives biofilm formation. Nat Commun 12:1399. 10.1038/s41467-021-21718-y

62. Klinman JP, Rose IA. 1971. Purification and kinetic properties of aconitate isomerase from Pseudomonas putida. Biochemistry 10:2253–2259. 10.1021/bi00788a011.

63. Thompson JF, Schaefer SC, Madison JT. 1990. Determination of aconitate isomerase in plants. Anal Biochem 184:39–47. 10.1016/0003-2697(90)90008-w.

64. Du C, Cao S, Shi X, Nie X, Zheng J, Deng Y, Ruan L, Peng D, Sun M. 2017. Genetic and Biochemical Characterization of a Gene Operon for trans-Aconitic Acid, a Novel Nematicide from Bacillus thuringiensis. J Biol Chem 292:3517–3530. 10.1074/jbc.M116.762666.

65. You X, Tian J, Zhang H, Guo Y, Yang J, Zhu C, Song M, Wang P, Liu Z, Cancilla J, Lu W, Glorieux C, Wen S, Du H, Huang P, Hu Y. 2021. Loss of mitochondrial aconitase promotes colorectal cancer progression via SCD1-mediated lipid remodeling. Mol Metab 48:101203. 10.1016/j.molmet.2021.101203.

66. Palmieri EM, McGinity C, Wink DA, McVicar DW. 2020. Nitric Oxide in Macrophage Immunometabolism: Hiding in Plain Sight. Metabolites 10:429. 10.3390/metabo10110429.

67. Nüse B, Holland T, Rauh M, Gerlach RG, Mattner J. 2023. L-arginine metabolism as pivotal interface of mutual host-microbe interactions in the gut. Gut Microbes 15:2222961. 10.1080/19490976.2023.2222961.

68. Guo J, Li X, Wang D, Guo Y, Cao T. 2019. Exploring metabolic biomarkers and regulation pathways of acute pancreatitis using ultra-performance liquid chromatography combined with a mass spectrometry-based metabolomics strategy. RSC Adv 9:12162–12173. 10.1039/c9ra02186h

69. Han M, Xie M, Han J, Yuan D, Yang T, Xie Y. 2018. Development and validation of a rapid, selective, and sensitive LC-MS/MS method for simultaneous determination of D- and L-amino acids in human serum: application to the study of hepatocellular carcinoma. Anal Bioanal Chem 410:2517–2531. 10.1007/s00216-018-0883-3.

